# LC-Based Metabolomics and Multivariate Analyses Reveal Metabolite Variations in Different Propagation Methods of *Eurycoma longifolia* Roots Harvested at Different Ages and Locations

**DOI:** 10.1101/2020.02.27.967786

**Authors:** Nur Nabilah Alias, Kamalrul Azlan Azizan, Norlia Basherudin, Nor Hasnida Hassan, Nazirah Abdullah

## Abstract

*Eurycoma longifolia* is a well-known medicinal plant with pharmacological effects and important bioactive compounds such as alkaloids and quassinoids. The suitable age of harvesting *E. longifolia* root for commercial use is relatively unknown and could influence the overall bioactive compounds present in the plant. In this study, non-targeted liquid chromatography mass spectrometer (LC-MS) and multivariate analyses (MVA) were performed to determine the chemical constituent of aqueous extract of 3-month-old *E. longifolia* hairy root (HR) culture, 1-, 4- and 11-year-old harvested *E. longifolia* roots. Unsupervised principle component analysis (PCA) and supervised partial least square discriminant analysis (PLSDA) were applied to evaluate metabolic similarities and differences in *E. longifolia* roots and hairy root in response to different harvesting age, locations and propagation methods. A total of 34 significant buckets with variable importance in projection (VIP) exceeding 1 were selected and identified. It was found that putatively identified quassinoids were significantly higher in 1-, 4- and 11-year-old *E. longifolia* roots whereas putative canthin-6-one alkaloids were abundantly present in 3-month-old *E. longifolia* hairy root (HR). These findings may be applicable to improve the harvesting age and increase the content of bioactive compounds in *E. longifolia* roots.

## Introduction

*Eurycoma longifolia* Jack, locally known as Tongkat Ali, is an herbaceous tree of the family Simaroubaceae, commonly found in Southeast Asian countries like Malaysia, Indonesia, Thailand, Myanmar, Cambodia and Laos. *E. longifolia*, particularly its roots is traditionally used for improvement of general health and libido. *E. longifolia* roots have been reported to exhibit various pharmacological and biological activities such as antimicrobial [1-4], anti-diabetic [5], antimalarial [6, 7], aphrodisiac activities [8] and cytotoxicity against cancerous cells [9-11]. The chemical profiling of *E. longifolia* root extract indicates alkaloids and quassinoids derivatives as the major bioactive compounds found in this plant [12, 28]. Quassinoids, particularly eurycomanone and their derivatives were reported to be abundant in *E. longifolia* roots and responsible for various pharmacological properties.

The increasing demands of *E. longifolia* root as herbal medicinal products have been impeded by the slow growth and late maturity of this plant. The suitable harvesting age of *E. longifolia* roots is unclear, but for commercial use, *E. longifolia* roots are usually harvested at the age of four years old [13]. Today, approximately 21,000 kg of *E. longifolia* were harvested to meet a demand of more than 54,000 kg per year [12]. As a result, this practice has led to excessive exploitation, extinction and adulterations in *E. longifolia* based products. Therefore, there is a need to standardize harvesting age to ensure sufficient supply and sustainability of this plant. In vitro production of *E. longifolia* such as micropropagation technique [14], callus culture [15, 16], cell suspension culture [17] and hairy root culture [18-20] have been proposed to preserve and supply the increasing demands of this plant. However, different propagation methods of *E. longifolia* root and harvesting age can influence the production of bioactive compounds, thus affecting the consistency of *E. longifolia* phytochemical content. In particular, the metabolic profiles and chemical differences in relation to different harvesting age of *E. longifolia* roots have not been elucidated systematically and is of great interest for improving the quality of *E. longifolia* roots.

Due to the chemical complexity of *E. longifolia* roots extract, an efficient analytical methodology to identify and quantify bioactive compounds in *E. longifolia* is of great interest. Liquid chromatography (LC) based metabolomics approach has been widely used to elucidate the comprehensive metabolic profiles of natural products in herbal plants such as *Panax ginseng* [21] and *Ginkgo biloba* L [22]. Liquid chromatography with mass spectrometry (LC-MS) is recognized as a powerful tool for identification and quantification of various herbal product and their constituents [23]. Previously, Chua et al. [24] reported a LC based metabolomics and multivariate analysis to discriminate *E. longifolia* roots from different locations and extraction temperatures. Similarly, Ebrahimi et al. [25] developed a nuclear magnetic resonance (NMR) based metabolomics approach to differentiate *E. longifolia* roots from different geographical locations, in relation to the biologically active quassinoids. In particular, both studies have discovered that quassinoids are major compounds in *E. longifolia* and thus, can be used as a chemical marker for the classification of *E. longifolia* roots.

The aim of this study was to investigate how different harvesting age, locations and propagation methods (seed and tissue culture) affected the chemical content of *E. longifolia*. Specifically, we performed untargeted LC-QTof-MS based metabolomics to determine metabolic contents of *E. longifolia* hairy root culture harvested at 3 months and roots of *E. longifolia* harvested at 1 year, 4 years and 11 years old. Multivariate analyses was used to find separation trends between different harvesting age and highlight metabolites that differentiated the *E. longifolia* roots and hairy root (HR). The results of this study would provide useful information on appropriate harvesting age in order to ensure the consistency of the bioactive compounds in *E. longifolia* roots for standardization purposes.

## Materials and methods

### Sample preparation

In this study, a total of 24 *E. longifolia* root samples belonging to four different harvested ages (3-month-old hairy root (HR) culture, 1-, 4- and 11-year-old roots) were subjected to metabolite extraction and LC-QToF-MS analysis. *E. longifolia* roots at the age of 4 and 11 years were collected from FRIM Research Sub-station Maran, Pahang (3.6383° N, 102.8052° E). 1-year-old *E. longifolia* samples were kindly provided by FRIM Ethnobotanical Garden (3.235339° N, 101.634269° E). 3-month-old *E. longifolia* hairy root cultures were supplied by FRIM Tissue Culture Laboratory. The *E. longifolia* of 1, 4 and 11 years old were previously propagated from seeds while the 3 months old *E. longifolia* hairy root was derived from successful transformation of *E. longifolia* target tissue using *Agrobacterium rhizogenes*. Upon harvesting, all the *E. longifolia* root samples were immediately placed in liquid nitrogen and stored at −80°C until extraction process.

### Metabolites extraction

All used solvents were purchased from Fisher Scientific (Loughborough, United Kingdom). Root samples from each harvested *E. longifolia* roots were chopped, freeze-dried with liquid nitrogen and ground to fine particles using mortar and pestle. 0.1 gram of the powdered root was extracted with solvent combinations, comprising 200 µL chloroform, 470 µL methanol and 80 µL water (20:47:8). All the extracts were centrifuged, filtered and analysed using LC-QTOF-MS. 50µL of internal standard (100ppm) was spiked into all samples before analysis.

### LC-QTOF-MS parameter

10 µL of *E. longifolia* extract was injected into a Dionex Ultimate 3000 UPLC-system (Thermo Scientific, USA) coupled to a micrOTOF-Q hybrid QTOF mass spectrometer (Bruker Daltonics, Bremen, Germany). Separation was carried out using an Acclaim RSLC 120 C18 (2.2 µm, 2.1 × 100 mm column) (Thermo Scientific, USA). The column was maintained at 40 °C and the flow rate was set to 0.2 mL/min. The mobile phases consisted of 0.1% formic acid (solvent A) and 100% acetonitrile (solvent B) with a total run of 22 min. The gradient began at 5% B (0 - 3 min), 80% B (3 - 10 min), 80% B (10 - 15 min) and 5% B (5 - 22 min). Eluted compounds were detected from 50 m/z to 1000 m/z at a scan accumulation rate of 2 scan/s and interscan delay of 0.1 s. MicrOTOF-Q was set with the following settings: nebulizer pressure, 1.2 bar; drying gas: 8 L/min at 200 °C, capillary voltage, 4500 V; end plate offset, - 500 V; funnel 1 RF, 200 Vpp; and funnel 2 RF, 200 Vpp. All samples were analyzed in positive ionization mode. The system was controlled by the HyStar software (version 3.2, Bruker) and micrOTOF control software (version 3.4, Bruker Daltonics, Bremen, Germany).

### Data pre-processing

Raw data from LC-QTOF-MS were exported into ProfileAnalysis version 2.1 (Bruker Daltonics, Bremen, Germany) for data alignment and bucketing. Find molecular features (FMF) was applied to search for peaks with the following parameters: bucketing using time alignment; S/N 5; correlation coefficient, 0.7; minimum compound length, 12 spectra; and smoothing width, 5. A filter was set (>30%) to remove bucket signals that had missing peaks (ion intensity=1) in the samples in any group. Subsequently, a dataset that contained buckets (RT-m/z pair) and their respective intensities (extracted ion chromatogram intensities) were obtained.

### Multivariate Analysis (MVA)

MetaboAnalyst 4.0 server (www.metaboanalyst.ca) was used for data normalization (by sum), log transformation, pareto scaling, statistical analysis and heatmap generation. Two-way analysis of variance (ANOVA) was performed in order to identify significantly different metabolites in different ages of *E. longifolia* roots. Significant differences between the means were determined at a 5% level of probability (p *<* 0.05). Buckets with significance (p < 0.05) were expressed as the mean ± standard error and subjected to a multivariate analysis using MetaboAnalyst 4.0 server. Unsupervised principal component analysis (PCA) was conducted to visualize the grouping patterns and to detect the outliers in the datasets. Supervised partial least square discriminant analysis (PLSDA) was performed to find a potential marker compound for each sample. The quality of the PCA and PLS-DA models was described by the cross-validation parameters, R^2^ and Q^2^, representing the explained variance and the predictive capability of the model, respectively. The candidate marker compounds were checked for their contribution to the discrimination trends by using variable important for projection (VIP) scores of PLSDA. The VIP score was also used to screen for highly important metabolites, whose significance was further verified through ANOVA. Buckets of interest were further identified using accurate mass and fragmentation patterns by comparing against published literature and publicly available online metabolite databases such as METLIN (https://metlin.scripps.edu), Massbank (http://www.massbank.jp) and Metfrag (http://msbi.ipb-halle.de/MetFrag/).

### Pathway Analysis

Pathway analysis was also conducted for the metabolites that significantly changed with p < 0.05 using MetaboAnalyst 4.0 software via Kyoto Encyclopedia of Genes and Genomes (KEGG) pathway database (http://www.genome.ad.jp/kegg/pathway.html) by comparing with *Arabidopsis thaliana* (thale cress) and *Oryza sativa japonica* (Japanese rice) pathway libraries. The mass error tolerance was < 10ppm.

## Results and Discussion

### Metabolite Profiling of *E. longifolia* Roots

Chemical profiles of *E. longifolia* hairy root and *E. longifolia* roots harvested at different maturity age were determined using LC-QToF-MS. Figure 1 shows the representative chromatograms of four *E. longifolia* roots, harvested at different age (3-month-old hairy root culture, 1-, 4- and 11-year-old roots). Clear differences in the chromatogram was observed for hairy root and 4 years old of *E. longifolia* roots when compared to those in 1 year and 11 years old of *E. longifolia* roots. Further analyses resulted in 162 buckets (RT-m/z pair) and subjected for multivariate analyses.

**Figure 1.**
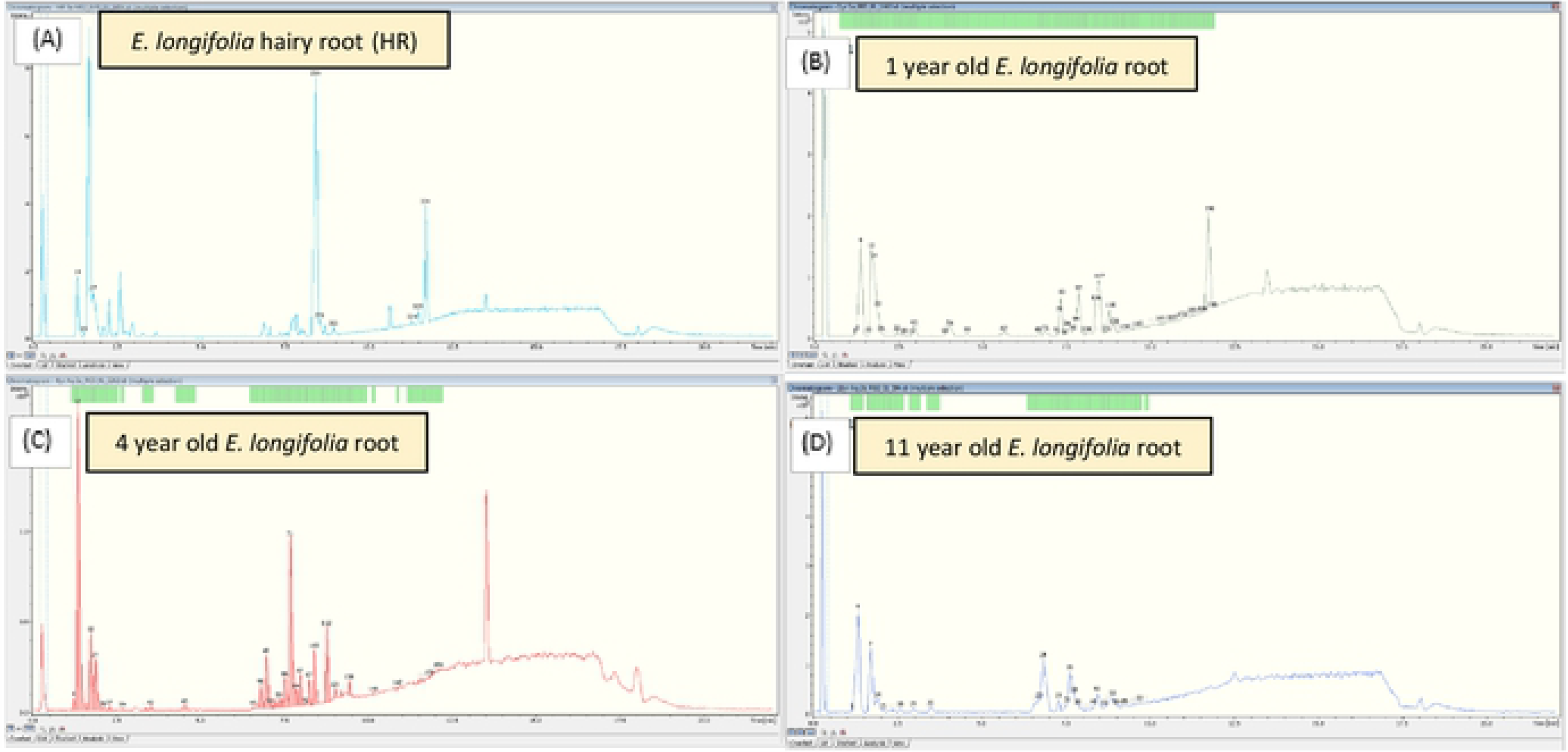
Representative LC-QTof-MS chromatogram of *E. longifolia* root harvested at different age (A) hairy root, (B) 1 year old, (C) 4 years old, (D) 11 years old *E*.*longifolia* root.

### Principal Component Analysis (PCA)

Unsupervised PCA was performed to visualize the separation trends between different *E. longifolia* samples. As showed in Figure 2, a clear separation between samples was obtained in the PCA score plot using the first two principal components (PC1 versus PC2) with total variance (R^2^) of 43.5% and Q^2^ (cum) of 11.9% (Figure 2A). The PCA score plot indicated that the hairy root (HR) culture (blue colour) was farther separated in the lower right quadrant of the score plot. This clearly showed strong variation between hairy root and mature roots of *E. longifolia*. Similarly, 1 year old *E. longifolia* root was separated away in the upper quadrant. This observation suggested a strong difference in the metabolite composition of *E. longifolia* roots in response to different harvesting age. Meanwhile, an overlapped between 11 years old (green colour) and 4 years old (purple colour) *E. longifolia* roots was observed in the lower left quadrant of PCA score plot. This indicated similarities in the metabolites levels from these two samples. The PCA scatter loading plot (Figure 2B) showed the projection of buckets that could contribute to the separation patterns in the PCA score plot.

**Figure 2.**
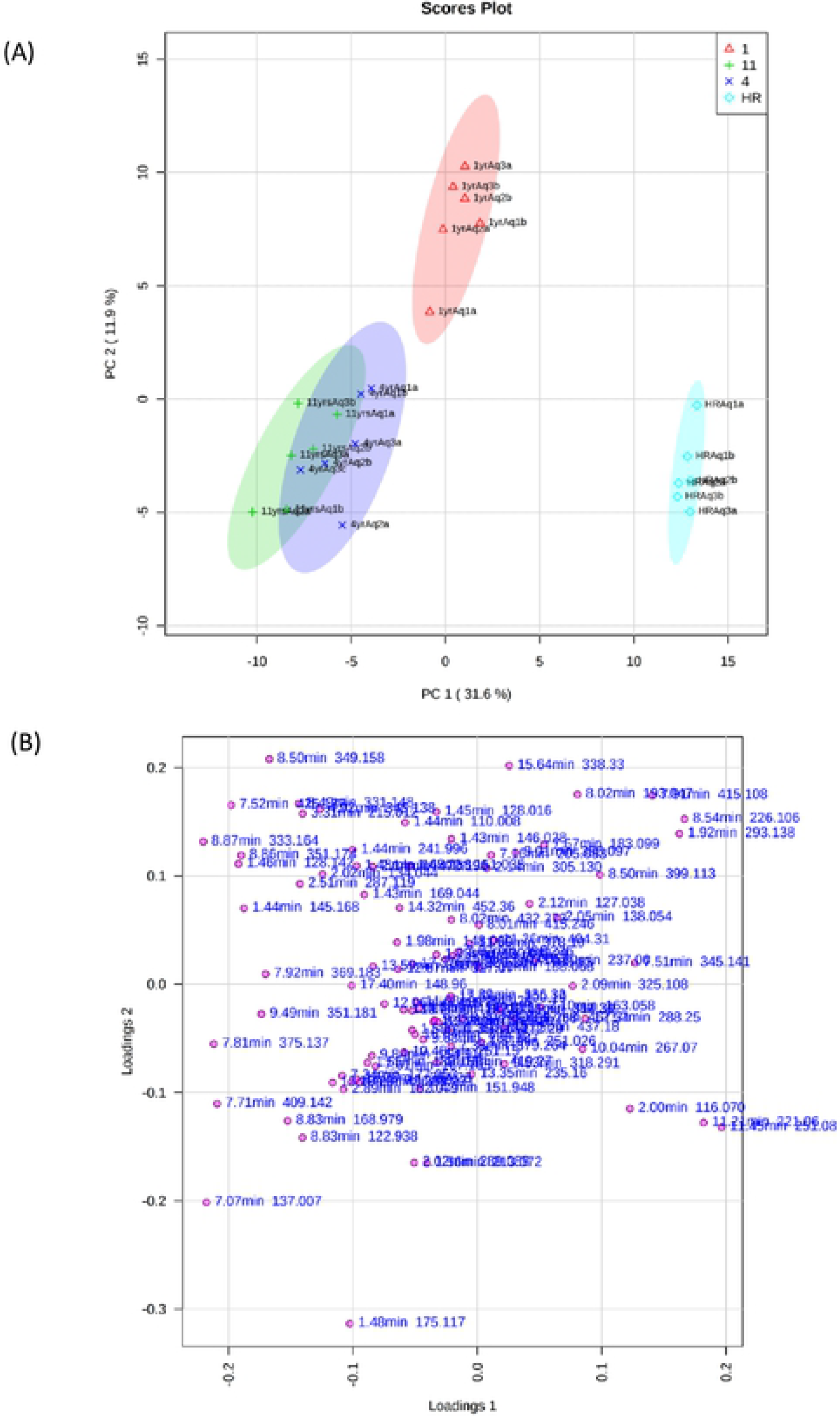
Principal component analysis (PCA) of buckets derived from 3-month-old hairy root culture (HR), 1, 4 and 11 years old of *E. longifolia* roots. (A) PCA score plot of PC1 versus PC2, (B) contribution of the buckets through the loadings of PCA scatter loading plot

### Partial Least Square Discriminant Analysis (PLSDA)

Supervised PLSDA was conducted to enhance the separation patterns obtained from PCA model and subsequently to identify potential marker compounds that contribute to the separation between tested *E. longifolia* hairy root and root samples. The PLSDA model required two latent variables to explain the discrimination trends between the *E. longifolia* hairy root and root samples. The values of total variation (R^2^X_(cum)_), model robustness (R^2^Y_(cum)_) and predictive ability (Q^2^_(cum)_) of the PLSDA model were used to further evaluate the quality of PLSDA models. The PLSDA score plot demonstrated better discrimination trends between *E. longifolia* hairy root and root samples at different ages with R^2^X_(cum)_ and Q^2^_(cum)_ value of 94.6% and 84.8%, respectively (Figure 3A). In common practise, R^2^X_(cum)_ and Q^2^_(cum)_ values exceeding 0.5 suggest a better fitting and more accurate model [26].

**Figure 3.**
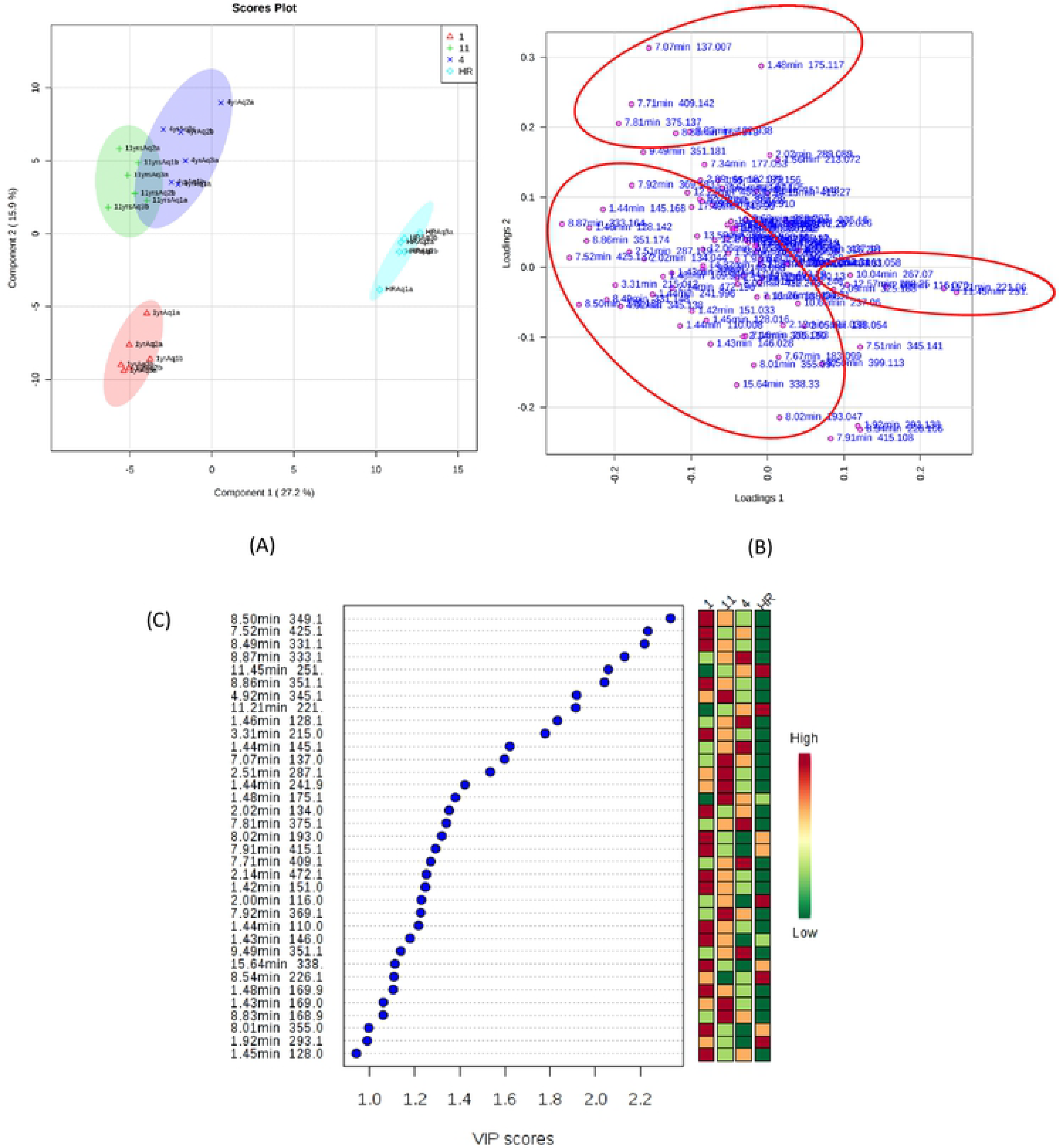
Partial least squares discriminant analysis (PLSDA) of buckets derived from 3 months old of hairy root culture (HR), 1, 4 and 11 years old of *E. longifolia* roots. (A) PLSDA score plot of PC1 versus PC2, (B) contribution of the buckets through the loadings of PLSDA scatter loading plot. (C) VIP values determine the most relevant variables responsible for the separation

Permutation test with 100 iterations and cross-validation predictive residual (CV-ANOVA) indicated that the constructed PLSDA model was valid and highly significant (*p* [CV-ANOVA] < 0.01). The loading plot corresponding with PLSDA score plot revealed signatory buckets that were important for the discrimination trends (Figure 3B). The majority of important marker compounds were located at upper, lower left and right quadrants of the scatter loading plot. In order to further identify buckets of interest in different *E. longifolia* roots, all buckets were screened based on the selection criteria of VIP value above 1 (VIP >1). A total of 34 buckets with VIP values exceeding 1 were discovered and were putatively identified (Figure 3C). Bucket belonging to 8.50min m/z 349.158 had the highest VIP value (2.2768) and was putatively identified as eurycomalactone (C_19_H_24_O_6_).

### Analysis of variance (ANOVA)

In addition to the VIP scores, buckets contributing to the discrimination trends were further validated using one-way ANOVA to evaluate their statistical significance. A total of 50 buckets were found to be statistically significant (*p* ≤ 0.05) (Table 1) whereas 12 buckets contributing to the discrimination trends in PLSDA model were putatively identified and listed in Table 2.

**Table 1.** A list of 50 compounds that are significant using Fisher’s Least Significant Difference (LSD) method (*p* ≤ 0.05)

| Compounds | t.value | p.value | $-\log_{10}(p)$ | FDR | Fisher's LSD |
| --- | --- | --- | --- | --- | --- |
| 1 7.07min 137.007m/z | 2119.80 | 0.00 | 24.47 | 0.00 | 11 - 1; 4 - 1; 11 - 4; 11 - HR; 4 - HR |
| 2 8.87min 333.164m/z | 1483.10 | 0.00 | 22.93 | 0.00 | 4 - 1; 1 - HR; 4 - 11; 11 - HR; 4 - HR |
| 3 11.45min 251.080m/z | 1139.20 | 0.00 | 21.79 | 0.00 | HR - 1; HR - 11; HR - 4 |
| 4 11.21min 221.069m/z | 691.97 | 0.00 | 19.64 | 0.00 | HR - 1; HR - 11; HR - 4 |
| 5 1.46min 128.142m/z | 366.76 | 0.00 | 16.92 | 0.00 | 4 - 1; 1 - HR; 4 - 11; 11 - HR; 4 - HR |
| 6 7.52min 425.137m/z | 317.63 | 0.00 | 16.31 | 0.00 | 1 - 11; 1 - 4; 1 - HR; 11 - HR; 4 - HR |
| 7 8.50min 349.158m/z | 43.68 | 0.00 | 8.24 | 0.00 | 1 - 4; 1 - HR; 11 - 4; 11 - HR; 4 - HR |
| 8 7.71min 409.142m/z | 38.31 | 0.00 | 7.75 | 0.00 | 11 - 1; 4 - 1; 1 - HR; 11 - HR; 4 - HR |
| 9 8.54min 226.106m/z | 33.87 | 0.00 | 7.31 | 0.00 | 1 - 11; 1 - 4; HR - 11; HR - 4 |
| 10 3.31min 215.012m/z | 33.19 | 0.00 | 7.23 | 0.00 | 1 - HR; 11 - HR; 4 - HR |
| 11 1.44min 145.168m/z | 33.10 | 0.00 | 7.22 | 0.00 | 1 - HR; 11 - HR; 4 - HR |
| 12 8.49min 331.148m/z | 27.65 | 0.00 | 6.59 | 0.00 | 1 - 4; 1 - HR; 11 - 4; 11 - HR; 4 - HR |
| 13 8.86min 351.174m/z | 27.20 | 0.00 | 6.53 | 0.00 | 1 - HR; 11 - HR; 4 - HR |
| 14 12.23min 359.236m/z | 25.66 | 0.00 | 6.33 | 0.00 | 11 - 1; 11 - 4; 11 - HR |
| 15 12.23min 354.281m/z | 25.60 | 0.00 | 6.33 | 0.00 | 11 - 1; 11 - 4; 11 - HR |
| 16 7.91min 415.108m/z | 25.24 | 0.00 | 6.28 | 0.00 | 1 - 11; 1 - 4; HR - 11; HR - 4 |
| 17 7.92min 369.183m/z | 24.43 | 0.00 | 6.17 | 0.00 | 11 - 1; 4 - 1; 1 - HR; 11 - HR; 4 - HR |
| 18 7.81min 375.137m/z | 22.50 | 0.00 | 5.89 | 0.00 | 11 - 1; 4 - 1; 1 - HR; 11 - HR; 4 - HR |
| 19 4.92min 345.138m/z | 20.58 | 0.00 | 5.60 | 0.00 | 1 - 4; 1 - HR; 11 - 4; 11 - HR; 4 - HR |
| 20 2.09min 325.108m/z | 19.62 | 0.00 | 5.45 | 0.00 | 1 - 11; HR - 1; 4 - 11; HR - 11; HR - 4 |
| 21 9.49min 351.181m/z | 19.50 | 0.00 | 5.43 | 0.00 | 11 - 1; 4 - 1; 1 - HR; 11 - HR; 4 - HR |
| 22 8.50min 399.113m/z | 15.39 | 0.00 | 4.70 | 0.00 | 1 - 11; 1 - 4; HR - 11; HR - 4 |
| 23 12.06min 415.206m/z | 14.85 | 0.00 | 4.59 | 0.00 | 11 - 1; 11 - 4; 11 - HR |
| 24 2.00min 116.070m/z | 14.76 | 0.00 | 4.58 | 0.00 | 11 - 1; HR - 1; 11 - 4; HR - 11; HR - 4 |
| 25 8.83min 372.971m/z | 13.11 | 0.00 | 4.23 | 0.00 | 11 - 1; 11 - 4; 11 - HR |
| 26 8.83min 168.979m/z | 12.80 | 0.00 | 4.17 | 0.00 | 11 - 1; 4 - 1; 11 - HR; 4 - HR |
| 27 8.83min 122.938m/z | 12.54 | 0.00 | 4.11 | 0.00 | 11 - 1; 4 - 1; 11 - HR; 4 - HR |
| 28 12.23min 436.357m/z | 11.12 | 0.00 | 3.78 | 0.00 | 11 - 1; 11 - 4; 11 - HR |
| 29 2.02min 134.044m/z | 10.06 | 0.00 | 3.52 | 0.00 | 1 - HR; 11 - HR; 4 - HR |
| 30 2.51min 287.119m/z | 9.84 | 0.00 | 3.47 | 0.00 | 1 - HR; 11 - HR; 4 - HR |
| 31 10.04min 267.074m/z | 8.31 | 0.00 | 3.06 | 0.00 | HR - 1; HR - 11; HR - 4 |
| 32 7.51min 345.141m/z | 7.94 | 0.00 | 2.95 | 0.00 | 1 - 11; 1 - 4; HR - 11; HR - 4 |
| 33 2.89min 182.079m/z | 7.55 | 0.00 | 2.84 | 0.00 | 11 - 1; 11 - 4; 11 - HR |
| 34 9.84min 154.910m/z | 7.32 | 0.00 | 2.77 | 0.00 | 11 - 1; 11 - 4; 11 - HR |
| 35 7.34min 177.053m/z | 7.28 | 0.00 | 2.76 | 0.00 | 11 - 1; 4 - 1; 11 - HR; 4 - HR |
| 36 8.02min 193.047m/z | 7.24 | 0.00 | 2.75 | 0.00 | 1 - 4; 11 - 4; HR - 4 |
| 37 1.44min 241.996m/z | 6.90 | 0.00 | 2.65 | 0.01 | 1 - HR; 11 - HR; 4 - HR |
| 38 1.48min 175.117m/z | 6.58 | 0.00 | 2.55 | 0.01 | 11 - 1; 4 - 1; 11 - HR |
| 39 1.43min 146.028m/z | 6.40 | 0.00 | 2.49 | 0.01 | 1 - 4; 1 - HR; 11 - 4; 11 - HR |
| 40 2.15min 130.085m/z | 6.34 | 0.00 | 2.47 | 0.01 | 1 - 4; 11 - 4; 11 - HR |
| 41 2.14min 472.196m/z | 5.91 | 0.00 | 2.33 | 0.01 | 1 - HR; 11 - HR; 4 - HR |
| 42 1.92min 293.138m/z | 5.60 | 0.01 | 2.23 | 0.01 | 1 - 4; HR - 11; HR - 4 |
| 43 10.48min 251.159m/z | 5.37 | 0.01 | 2.15 | 0.02 | 11 - 1; 11 - 4; 11 - HR |
| 44 1.48min 169.958m/z | 5.18 | 0.01 | 2.08 | 0.02 | 1 - HR; 11 - HR; 4 - HR |
| 45 1.44min 110.008m/z | 4.45 | 0.01 | 1.82 | 0.03 | 1 - 4; 1 - HR; 11 - HR |
| 46 1.55min 189.156m/z | 4.18 | 0.02 | 1.72 | 0.04 | 4 - 1; 11 - HR; 4 - HR |
| 47 1.45min 128.016m/z | 4.15 | 0.02 | 1.71 | 0.04 | 1 - 11; 1 - HR; 4 - HR |
| 48 1.42min 151.033m/z | 4.08 | 0.02 | 1.69 | 0.04 | 1 - 4; 1 - HR; 11 - HR |
| 49 14.15min 419.272m/z | 4.06 | 0.02 | 1.68 | 0.04 | 4 - 1; 4 - 11; 4 - HR |
| 50 17.40min 148.968m/z | 3.89 | 0.02 | 1.61 | 0.05 | 11 - HR; 4 - HR |

**Table 2.** Composition of quassinoids and cantin-6-one alkaloids in the aqueous solvent of *E. longifolia* extract using LC-MS

| Bucket | Class | Putative identification | Precursor ion | *p* value | VIP value | Molecular Formula | Predicted m/z | Measured m/z | References |
| --- | --- | --- | --- | --- | --- | --- | --- | --- | --- |
| **8.87 min – m/z 333.164** | Quassinoids | Eurycolactone C* | [M+H]<sup>+</sup> | <0.01 | 2.1304 | C<sub>18</sub>H<sub>26</sub>O<sub>6</sub> | 333.126 | 333.164 | [24] |
| **8.50 min – m/z 349.158** | Quassinoids | Eurycomalactone* | [M+H]<sup>+</sup> | <0.01 | 2.3341 | C<sub>19</sub>H<sub>24</sub>O<sub>6</sub> | 348.157 | 349.158 | [24], [27] |
| **8.86 min – m/z 351.174** | Quassinoids | 7α-Hydroxyeurycomalactone* | [M+H]<sup>+</sup> | <0.01 | 2.0412 | C<sub>19</sub>H<sub>26</sub>O<sub>6</sub> | 350.173 | 351.174 | [24] |
| **8.86 min – m/z 351.174** | Quassinoids | Eurycolactone E* | [M+H]<sup>+</sup> | <0.01 | 2.0412 | C<sub>19</sub>H<sub>26</sub>O<sub>6</sub> | 350.173 | 351.174 | [24] |
| **8.86 min – m/z 351.174** | Quassinoids | Eurycomalide A* | [M+H]<sup>+</sup> | <0.01 | 2.0412 | C<sub>19</sub>H<sub>26</sub>O<sub>6</sub> | 350.173 | 351.174 | [24] |
| **7.92 min – m/z 369.183** | Quassinoids | 2,3-Dehydro-4α-hydroxylongilactone* | [M+H]<sup>+</sup> | <0.01 | 1.2263 | C<sub>19</sub>H<sub>28</sub>O<sub>7</sub> | 368.184 | 369.183 | [24] |
| **7.71 min – m/z 409.142** | Quassinoids | Eurycomanone* | [M+H]<sup>+</sup> | <0.01 | 1.2707 | C<sub>20</sub>H<sub>24</sub>O<sub>9</sub> | 408.142 | 409.142 | [27] |
| **7.52 min – m/z 425.137** | Quassinoids | 13α-(21)-epoxyeurycomanone* | [M+H]<sup>+</sup> | <0.01 | 2.2333 | C<sub>20</sub>H<sub>24</sub>O<sub>10</sub> | 424.137 | 425.137 | [24], [27] |
| **11.21 min – m/z 221.069** | Canthin-6-one alkaloids | Canthin-6-one* | [M+H]<sup>+</sup> | <0.01 | 1.9139 | C<sub>14</sub>H<sub>8</sub>N<sub>2</sub>O | 220.064 | 221.069 | [24] |
| **11.45 min – m/z 251.080** | Canthin-6-one alkaloids | 9-methoxycanthin 6-one* | [M+H]<sup>+</sup> | <0.01 | 2.0582 | C<sub>15</sub>H<sub>10</sub>N<sub>2</sub>O<sub>2</sub> | 250.074 | 251.080 | [24], [28] |
| **10.04 min – m/z 267.074** | Canthin-6-one alkaloids | 1-Hydroxy-11-methoxycanthin-6-one | [M+H]<sup>+</sup> | <0.01 | 0.9171 | C<sub>15</sub>H<sub>10</sub>N<sub>2</sub>O<sub>3</sub> | 266.069 | 267.074 | [24] |
| **8.50 min – m/z 399.113** | Canthin-6-one alkaloids | 11-o-β-D-Glucopyranosylcanthin-6-one | [M+H]<sup>+</sup> | <0.01 | 0.5814 | C<sub>20</sub>H<sub>18</sub>N<sub>2</sub>O<sub>7</sub> | 398.111 | 399.113 | [24] |

\* - highly influenced metabolites (VIP ≥ 1.00 and *p* ≤ 0.05)

Three important buckets belonging to quassinoids; 8.50 min – m/z 349.158 putatively identified as eurycomalactone (C_19_H_24_O_6_) [24, 27], 7.71 min – m/z 409.142m/z putatively assigned as eurycomanone (C_20_H_24_O_9_) [27] and 7.52 min – m/z 425.137 putatively identified as 13α-(21)-epoxyeurycomanone [24, 27] were detected. Eurycomanone is the most abundant phytochemical quassinoids in *E. longifolia* roots [24, 27 and 29] which has stable composition (not easily metabolized) [30]. A study by Low et al. [31] reported that eurycomanone and 13α-(21)-epoxyeurycomanone possessed good physicochemical properties such as chemical stability, plasma stability, plasma protein binding, solubility and permeability. In fact, both of them are the main quassinoids contributing to the overall antimalarial activity of *E. longifolia*. Also, these two compounds including eurycomalactone are usually used as standard markers for the standardization of *E. longifolia* products [24].

Concomitantly, buckets belonging to canthin-6-one alkaloids were 11.45 min-m/z 251.080 putatively identified as 9-methoxycanthin 6-one (C_15_H_10_N_2_O_2_) [24, 28], 11.21 min – m/z 221.0.69 assigned to canthin-6-one (C_14_H_8_N_2_O) [24], 8.50 min – m/z 399.113 identified as 11-o-β-D-Glucopyranosylcanthin-6-one (C_15_H_10_N_2_O_3_) [24] and 10.04 min – m/z 267.074 putatively identified as 1-Hydroxy-11-methoxycanthin-6-one (C_15_H_10_N_2_O_3_) [24]. Canthin-6- one alkaloids are naturally occurring amine compound which is useful as insects and herbivores repellent [24]. Finding by Kuo et al. [32] demonstrated that canthin-6-one and 9- methoxycanthin 6-one exhibit a significant cytotoxicity effect against human lung cancer (A- 549) and human breast cancer (MCF-7) cell lines. Additionally, 9-methoxycanthin 6-one is also regarded as one of the standard markers for the standardization of *E. longifolia* products.

### Metabolite variations in *E. longifolia* hairy root (HR) and roots harvested at different ages

Figure 4 illustrated the distribution and relative quantification of 50 buckets with *p* ≤ 0.05 according to *E. longifolia* samples. The red and green colours in the heatmap represent higher and lower relative concentrations of selected buckets, respectively. One-way hierarchical cluster analysis showed buckets were grouped into two main clusters. Cluster 1 comprised buckets belonging to canthin-6-one alkaloids. Specifically, buckets assigned to 9- methoxycanthin 6-one, canthin-6-one, 11-o-β-D-glucopyranosylcanthin-6-one and 1-hydroxy-11-methoxycanthin-6-one were abundant in the hairy root (HR) but were lower in roots of 4 and 11 years old. The findings are similar to those reported by Tran et al. [20] that indicates the increment of 9-methoxycanthin 6-one in transgenic hairy root (HR) than those in the wild roots of *E. longifolia*.

**Figure 4.**
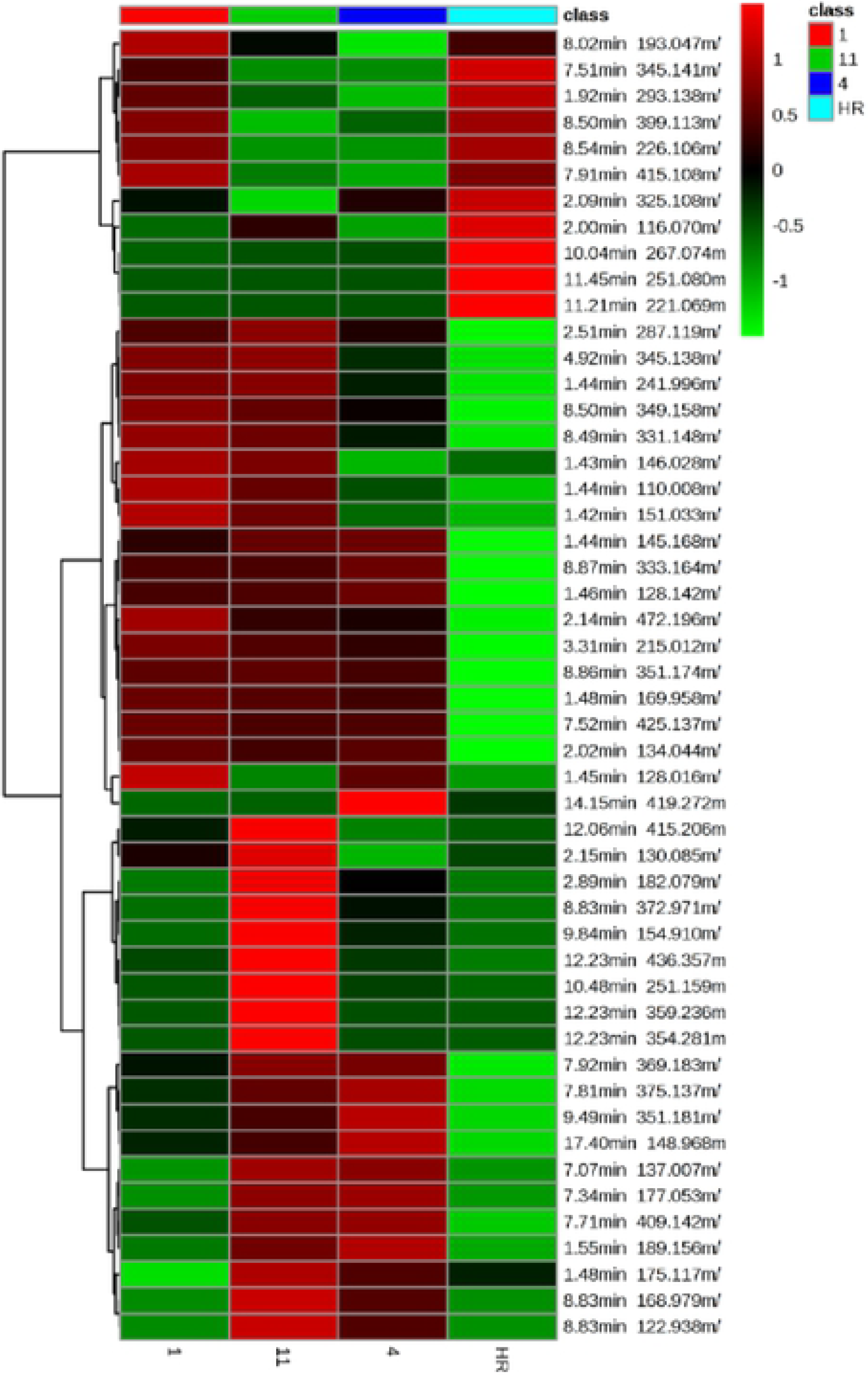
Top 50 buckets with significance and their relative levels in 3-month-old *E*. *longifolia* hairy root (HR) culture, 1-, 4- and 11 year-old *E. longifolia* roots. One-way hierarchical clustering heatmap was generated using MetaboAnalyst 4.0 server.

Cluster 2 was dominated by buckets that were putatively identified as quassinoids; eurycomanone, eurycomalactone, 13α-(21)-epoxyeurycomanone, eurycolactone C, eurycolactone E, eurycomalide A, 7α-hydroxyeurycomalactone and 2,3-dehydro-4α-hydroxylongilactone. These compounds were found higher in 4 and 11 years old *E. longifolia* roots but were lower in hairy root (HR) and 1 year old roots. Notably, both eurycomanone and eurycomalactone were abundant in 11 years old root. Furthermore, eurycomanone was also present in 4 years old root whereas eurycomalactone was highly expressed in 1 year old sample. In contrast, 13α(21)-epoxyeurycomanone, eurycolactone C, eurycolactone E, eurycomalide A, 7α-hydroxyeurycomalactone were abundantly found in *E. longifolia* roots of 1, 4 and 11 years old. The results suggested the accumulation of quassinoids in response to harvesting age. Also, it was found that quassinoids like eurycomanone might not be present in the roots at early age of the plant. Previous study by Nor-Hasnida et al. [14] highlighted that there was a difference of eurycomanone content between tissue culture plantlets and matured root of *E. longifolia*.

The findings in this current study indicated that the age of plant plays important role in the formation of secondary metabolites in *E. longifolia*, particularly eurycomanone. In fact, the results support the previous studies which hypothesized that *E. longifolia* can be harvested as early as 4-year-old for commercial use. Nevertheless, we are still lacking in evidence to confirm whether the presence of eurycomanone would be the best indicator in selecting high quality planting material of *E. longifolia* for commercialization purpose.

### The effect of different locations and propagation methods on chemical profile of *E. longifolia* roots

The effect of locations and propagation methods on chemical profile of *E. longifolia* root was assessed using PCA (Figure 5). Three clusters were observed (green, red and purple), demonstrating distinct metabolite variations in *E. longifolia* roots in response to different locations, harvesting age and propagation methods. It was observed that samples of 4 and 11 years old were overlapped (green colour), but were differentiated from 1 year old *E. longifolia* roots. This observation strongly suggested the effect of locations on the overall chemical constituent of *E. longifolia* roots. Meanwhile, hairy root (HR) culture was farther separated, indicating the effect of propagation methods on the chemical constituent of *E. longifolia*. As overall, the results proposed that location factors and propagation methods greatly affect the chemical profile of *E. longifolia* [33].

**Figure 5.**
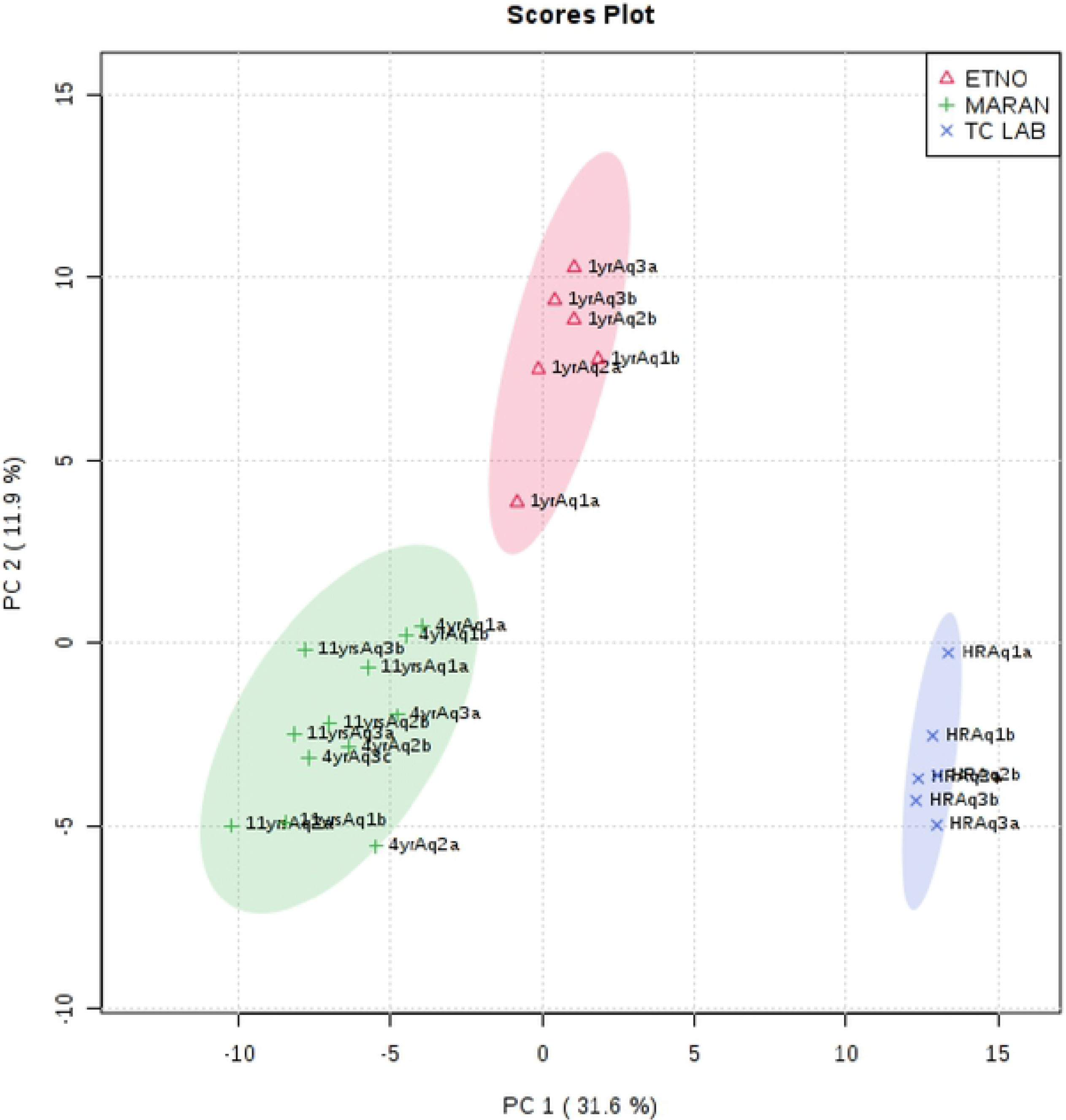
PCA score plot based on propagation methods and locations of the *E. longifolia*. ETNO = FRIM Ethnobotanical Garden; TC LAB = FRIM Tissue Culture Laboratory.

### Pathway analysis

Although there was a difference in many metabolite levels in *E. longifolia* roots in response to different locations, harvesting age and propagation methods, further analyses focused on the metabolites that exhibited the greatest differences between the samples. According to the results of PLSDA with the screening condition of VIP > 1 and p ≤ 0.05, 34 buckets were selected for the pathway analysis of which 12 of them were involved in various biosynthesis pathways such as terpenoid backbone biosynthesis, galactose metabolism, diterpenoid biosynthesis, ubiquinone and other terpenoid-quinone biosynthesis, purine metabolism, biosynthesis of secondary metabolites, biosynthesis of antibiotics, cysteine and methionine metabolism, amino sugar and nucleotide sugar metabolism, riboflavin metabolism, steroid hormone biosynthesis, nicotinate and nicotinamide metabolism, carotenoid biosynthesis, phenylalanine metabolism, histidine metabolism and many more (Table 3). Kuo et al. [32] believed that all quassinoids, for instance eurycomanone are biosynthesized through the triterpenoid biosynthesis pathway, where the process begins with the degradation of triterpenes.

**Table 3.**
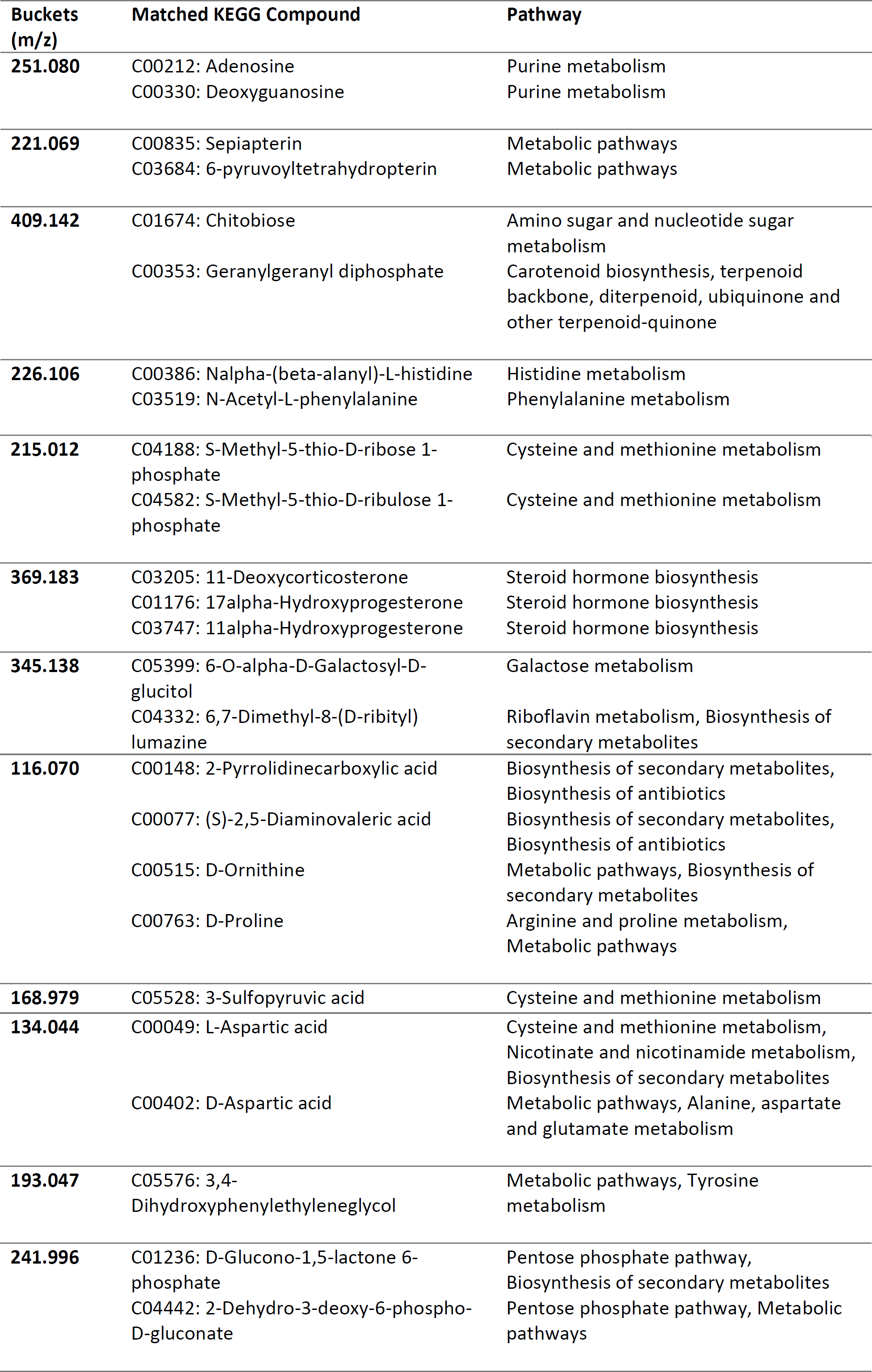
Buckets that match with KEGG compound and pathway

## Conclusion

In this study, non-targeted liquid chromatography based metabolomics was used to understand the chemical constituents of *E. longifolia* hairy root (HR) and *E. longifolia* roots harvested at different age. PCA and PLSDA clearly showed the effect of harvesting age, locations and propagation methods on chemical profile of *E. longifolia* roots. A total of 34 significant (P<0.05) buckets with VIP exceeding 1 were selected and putatively identified. Putatively identified quassinoids were abundantly presented in *E. longifolia* roots of 1, 4 and 11 years old whereas putative canthin-6-one alkaloids related compounds were higher in hairy root (HR) culture. These findings extended our knowledge that the harvesting age, locations and propagation methods of *E. longifolia* roots play important role in selecting high quality of *E. longifolia* roots for commercialization purpose.

## Acknowledgements

This research was funded by Forest Research Institute Malaysia (FRIM) under RPP grant. We thank Puan Sarah Ibrahim from Institute of Systems Biology (INBIOSIS) for her technical assistance in dealing LC-QTof-MS and members of Stesen Penyelidikan FRIM Maran (SMRN) especially Puan Hada Masayu Ismail @ Dahlan for the information given on sampling location. Also, special thanks to members of FRIM Genetics Laboratory for their technical support.

